# Secondary-structure prediction revisited: Pβ and Pc represent structures of amyloids and aid elucidating phenomena in interspecies transmissions of prion

**DOI:** 10.1101/073668

**Authors:** Yuzuru Taguchi, Noriyuki Nishida

**Affiliations:** Division of Cellular and Molecular Biology, Department of Molecular Microbiology and Immunology, Nagasaki University, Graduate School of Biomedical Sciences Basic Medical Science Building (8 floor), 1-12-4 Sakamoto, Nagasaki 852-8523, JAPAN, /

**Keywords:** Secondary structure prediction, prion, amyloid, β-sheet propensity, strain, species barrier

## Abstract

Prion is a unique infectious agent which consists solely of abnormally-folded prion protein (PrP^Sc^) but possesses virus-like features, e.g. existence of strain diversity, adaptation to new hosts and evolutionary changes. These biological phenomena were attributed to the structural properties of PrP^Sc^ due to lack of genetic material of prion. Therefore, regardless of incompatibility with high-resolution structural analysis, many structural models of PrP^Sc^ have been hypothesized based on limited structural information and, recently, models consisting solely of β-sheets and intervening loops/kinks have been suggested, i.e. parallel in-register β-sheet models and β-solenoid model. Given the relatively simple structural models of PrP^Sc^, we utilized values of theoretical β-sheet or random-coil propensity (Pβ or Pc, respectively) calculated by secondary structure prediction with a neural network to analyze interspecies transmissions of prion, because numerical conversion of the primary structures would enable quantitative comparison between PrP with distinct primary structures. Reviewing experiments in the literature, we ascertained biological relevance of Pβ and Pc and demonstrated how those parameters could aid interpretation and explain phenomena in interspecies transmissions. Our approach can lead to development of a versatile tool for investigation of not only prion but also other amyloids.

---

Today, many incurable neurodegenerative diseases including Alzheimer's disease, Parkinson's disease and tauopathy are known to be caused by misfolding and amyloid formation of constitutively-expressed proteins. Although the amyloidogenic proteins physiologically exist as monomers in normal conformations, once they are misfolded and form amyloid nuclei/fibers, the nuclei induce misfolding of the normal conformers and incorporate them into the amyloids to grow in size and number, a process referred to as seeding. Thus amyloids propagate and eventually cause diseases. Transmissible spongiform encephalopathies (TSEs) are also a group of diseases caused by misfolding of prion protein (PrP), including Creutzfeldt-Jakob disease (CJD) in humans, bovine spongiform encephalopathies (BSE) in cattle, chronic wasting disease (CWD) in elk and scrapie in sheep. Unlike other amyloids, propagation of the misfolded PrP (PrP^Sc^) is very efficient and behaves like a virus despite lack of nucleotides as an infectious agent called prion. For instance, a prion has preferred species for the host, e.g. BSE's preference for cattle or sheep, and transmissions to the other species are often inefficient with longer incubation periods, i.e. species barrier. However, once the barrier is overcome, the prion changes the host range and the secondary passage is more efficient with shorter incubations [1]. Moreover, in transmissions of TSE, clinicopathological traits of the donor, e.g. clinical courses, distribution of lesions and PrP deposition patterns, are faithfully reproduced in the recipient: mistaken as a virus, each prion with unique clinicopathological properties was called “strain". The protein-only hypothesis argues that strain-specific pathogenic information is enciphered in structures of PrP^Sc^ and is inherited when host-encoded normal conformer PrP (PrP^C^) is refolded by PrP^Sc^ in template-directed manner [1]. Hence, the structures of PrP^Sc^ are important, but remain undetermined due to its incompatibility with conventional high-resolution structure analysis. Regardless, structural information of PrP^Sc^ have been collected through various methods like electronmicroscopic analysis of scrapie fibrils [2], hydrogen/deuterium-exchange mass spectrometry [3], disulfide cross-links of *in vitro*-formed PrP fibrils [4] and *in silico* analysis [5,6] to create structural models of PrP^Sc^. Although models with remaining α-helices were previously postulated [2,4,5], recently models devoid of α-helix, e.g. the “β-solenoid model” [7] and the “parallel in-register intermolecular β-sheet model” [8–10] have been postulated as the more plausible models. However, so far, none of the models seems to elucidate the mechanisms underlying the virus-like properties of prions.

If PrP^Sc^ consists solely of β-sheets and intervening loops/kinks, the PrP^C^-PrP^Sc^ interactions would be also relatively simple, especially in the case of parallel in-register models where each region of the substrate PrP^C^ interacts with the corresponding region of the template PrP^Sc^. Given the simple interaction modality, we thought that numerical conversion of the primary structure of PrP by algorithms which render structural information of PrP^Sc^ enable quantitative comparison between PrP of different primary structures, facilitating interpretation of results of interspecies transmission experiments; besides, arithmetic operation of the converted values might reflect certain aspects of the PrP^C^-PrP^Sc^ interactions. Therefore, for this purpose we attempted to use theoretical propensity to β-sheet formation (Pβ) calculated by a neural-network secondary structure prediction (http://cib.cf.ocha.ac.jp/bitool/MIX/) [11], based on the following four assumptions: (i) PrP^Sc^ consists of only β-sheets and intervening loops/kinks; (ii) regional structures of PrP^Sc^ including positions of the β-strands are determined by 'actual' intrinsic propensity to β-sheet formation; (iii) since there is supposedly no α-helix in PrP^Sc^, theoretical propensity to α-helix (Pα) is ignored; and (iv) the Pβ value at each residue substitute for the actual β-sheet propensity and, whether large or small, represents the regional structures of PrP^Sc^. Those assumptions allow using Pβ-graph as a surrogate model of the structure of PrP^Sc^.

Here, we confirmed biological relevance of Pβ by application to *in vitro* experiments from the literature, and incidentally discovered that the propensity to random coil structure (Pc) values are also important, being apparently related to cross-seeding efficiencies of amyloidogenic peptides. Then, we reviewed the results of interspecies transmission experiments in the literature from the view point of the above assumptions. Pβ analysis visually aided interpretation of experimental transmissions involving heterologous PrP molecules and gave possible explanations to the varied incubations among different hamster species and the changes of host ranges after interspecies transmissions.

## Results and Discussion

### Pβ-graphs of representative species

In the beginning, we compared Pβ-graphs of PrP from representative species. They were mostly the same as expected from the high homology in the primary structures, except for variations in the heights of some peaks (Fig. 1B). For example, Syrian hamster had a relatively low peak centered at 110 (Fig. 1B, **blue arrow**), while human or elk PrP had outstandingly high peaks centered at the residue 140 or 175, respectively (Fig. 1B, **red or green arrow**). Significance of the differences in the heights of those Pβ peaks is shown in the following sections. Incidentally, we noticed that there are apparently more Pβ peaks in PrP than in Aβ42 or α-synuclein (Fig. 1C and D). According to the initial assumptions, the Pβ-peaks could be in β-sheets in the template PrP^Sc^ and the similar structures are induced in the substrate PrP^C^ through direct contacts, with the high-Pβ regions functioning as the interaction “interfaces” with the PrP^Sc^: In the case, the existence of many peaks might be responsible for the strain diversity of prions, as we previously hypothesized [12]. Whether the Pβ-peaks actually correspond to the regions which have high affinity for PrP^Sc^, i.e. candidates for the interfaces, was critical for the purpose and we attempted confirmation of biological relevance of Pβ values.

**Figure 1.**
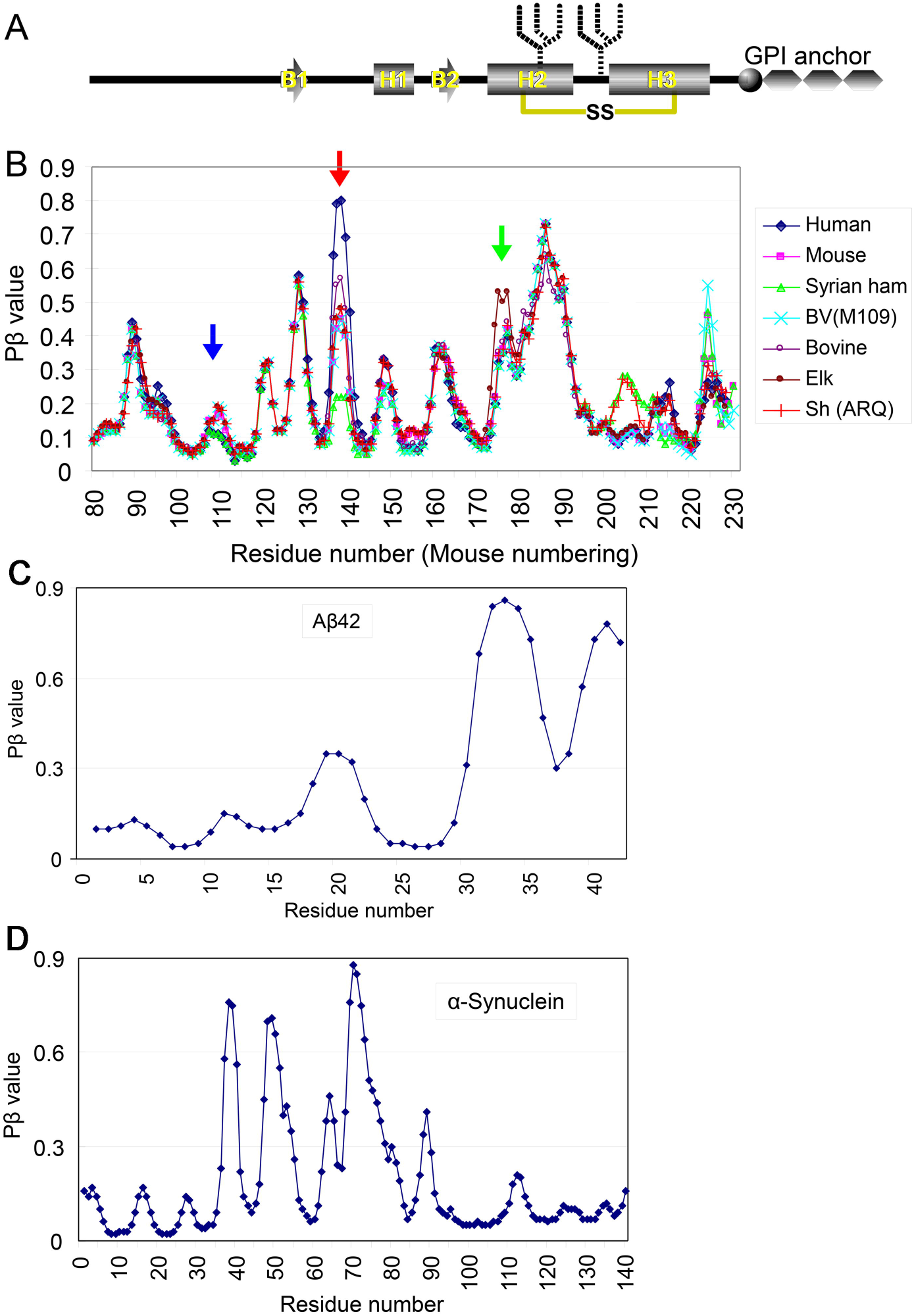
Pβ-graphs of PrP of representative species look very alike but with certain differences. **A.** A schematic illustration of the secondary-structure components of prion protein (PrP) and post-translational modifications, i.e. a disulfide link (yellow line with “SS”), N-linked glycans (fork-like objects) and glycosylphosphatidylinositol anchor (GPI anchor). B1 and B2, the first and the second -strands, respectively. H1, H2 and H3, the first, second and the third a-helix, respectively. **B.** Pβ-graphs of PrP from various species. The blue, red and green arrows indicate peaks centered at the residue 110, 140 and 175, respectively. Syrian ham, Syrian hamster. BV(109M), bank vole with methionine at the residue 109 (in bank-vole numbering). Sh(ARQ), sheep with A136/R150/Q171 polymorphism. **C and D.** Pβ-graphs of A 42 and human a-synuclein, respectively.

### Testing biological relevance: Application to ΔPrP-series and C-terminally-truncated mutant PrP

To test the biological relevance of the Pβ-graphs, we applied the algorithm to ΔPrP-series (Fig. 2A, **left panel**), which is a series of mutant PrP with various lengths of internal deletions in the region from the residue 159 to 175 [13]. Inverse correlation between the deletion sizes and efficiencies of dominant-negative inhibition (DNI) of the deletion mutants (Fig. 2A, **right panel**) implied that the region is the main interface for ΔPrP to bind the template PrP^Sc^. On the Pβ-graph of wild-type mouse PrP, two peaks, i.e. interface candidates, were recognized in the region 159-175 (Fig. 2B). As the deletion was elongated from the residue 159 to the C-terminal direction, the peak of Pβ centered at ~160 was gradually lowered (Fig. 2B or 2C, **red arrow**) and almost disappeared in Δ159-166. The other peak centered at ~175 (Fig. 2D, **black arrow**) also became narrower in Δ159-171 (Fig. 2D, **red arrow**) and finally disappeared in Δ159-175 (**blue line with open square**). Collectively, the gradual disappearances of the two peaks of Pβ-graphs of ΔPrP series seemed to correlate with the gradual decrease of DNI of the mutants [13]. The correlation of the alterations of Pβ-graph with the biological events substantiated a certain degree of its biological relevance.

**Figure 2.**
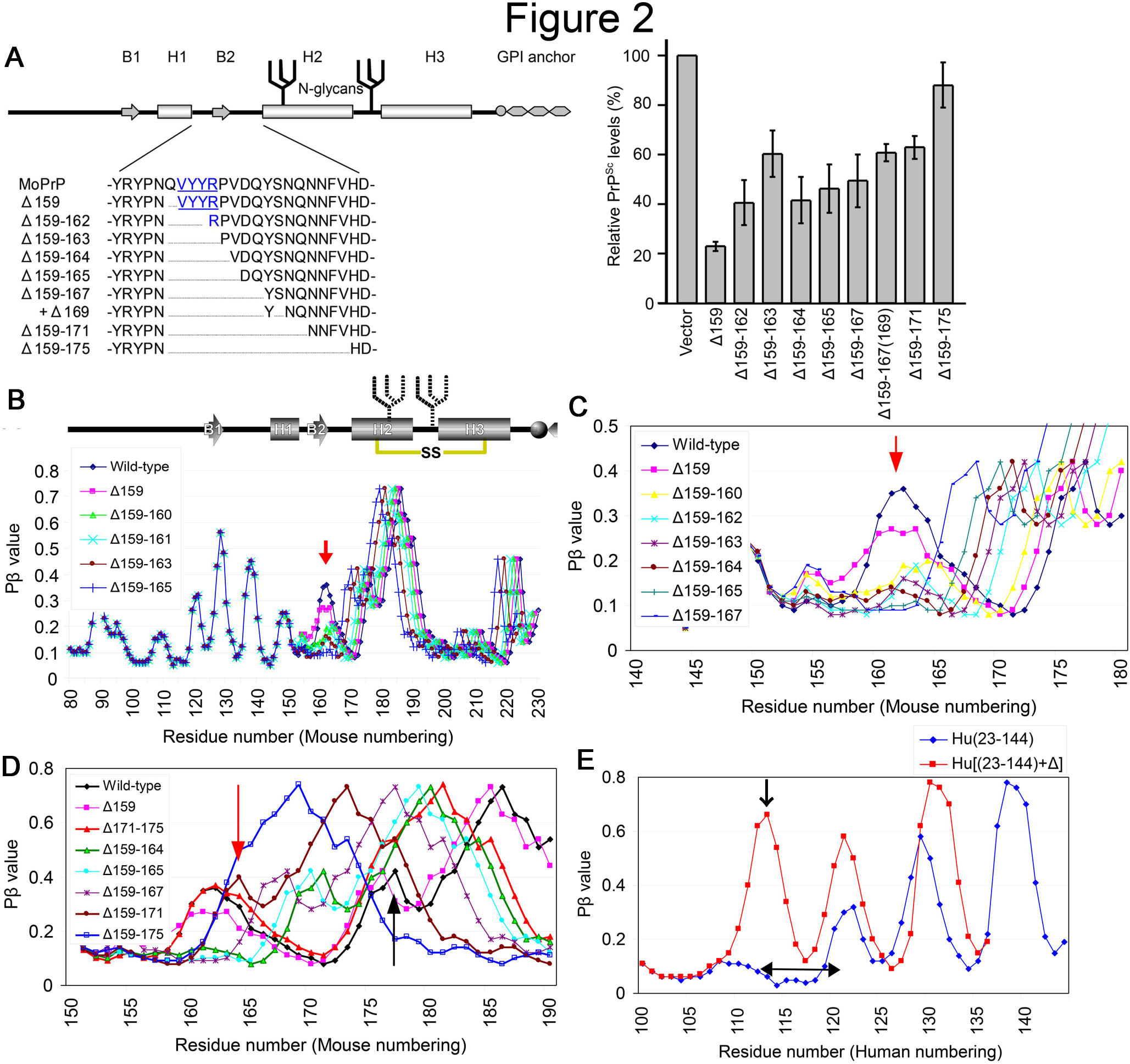
Biological relevance of Pf3: heights of Pf3 peaks correlate with dominant-negative inhibition of PrP-series mutants. **(Left panel)** A schematic illustration of the deletion-mutant PrP with deletions in the region between H1 and H2 PrP-series) and **(right panel)** a graph showing the PrPsc levels of convertible PrP coexisting with the indicated non­: onvertible PrP mutant on 22L-infected N2a cells, cited from [13]. Note that PrP^8^c levels of the convertible PrP ncrease, as the deletion size of the co-existing PrP enlarged: this means that the dominant-negative inhibitory ffects of the non-convertible PrP on the co-existng convertible PrP are decreased as the deletion elongates. v1oPrP, wild-type mouse PrP. **:l.** Pβ-graphs of wild-type mouse PrP and PrP mutants. Note that height of the peak at around the residue 160 is lradually lowered as the deletion is elongated (red arrow), while the appearances of the rest of the Pβ-graphs are 1ardly affected.:. Pβ-graphs of wild-type mouse PrP and PrP mutants, focused on the region from the residue 140 to 180, to show he diminution of the peak clearly.). Pβ-graphs of wild-type mouse PrP and PrP mutants focused on the region from the residue 150 to 190, focusing m another peak at around the residue 175 (black arrow). Note that the peak becomes narrower in the159-171 (red mow) and finally disappear in 159-175 (blue curve with open square).:. Pβ-graphs of C-terminally-truncated human PrP [Hu(23-144)] and a deletion variant of the truncated mutant Hu(23-144)+] which has internal deletion from the residue 113 to 120 (left-right arrow) [14]. Note that another large >eak appears by the deletion 113-120 (arrow).

We also applied the algorithm to the C-terminally-truncated mutant human PrP, PrP23-144. Jones and colleagues reported that a deletion of the region from the residue 113 to 120, which supposedly constitute a β-sheet core of the amyloid, did not affect amyloidogenicity of the deletion variant, to their surprise, and solid-state NMR of the variant revealed an altered β-core encompassing 106 to 125 [14]. On Pβ-graphs of PrP23-144 and the deletion variant (Fig. 2E), a large peak was newly generated by the internal deletion Δ113-120 albeit with discrepancy in its exact position from the result of solid-state NMR, suggesting a possibility that Pβ analysis could have predicted the newly-formed β-sheet core.

### Testing biological relevance: Significance of Pβ and Pc values on cross-seeding efficiencies

Regarding this type of truncated mutants, we were also interested in experiments of cross-seeding reaction among the PrP23-144 from human, mouse and Syrian hamster PrP [Hu(23-144), Mo(23-143) and Sy(23-144), respectively] [15]. The Hu(23-144) could be cross-seeded by *in vitro*-formed fibrils of Mo(23-V. but not by fibril of Sy(23-144), while Mo(P23-143) was cross-seeded by either fibril of Hu or Sy(23-144). In contrast, neither human nor mouse fibrils could seed Sy(23-144). Pβ-graphs of those molecules were clearly different in the C-terminal region, with the human outstandingly high, the hamster the lowest and the mouse at the intermediate (Fig. 3A): those differences seemed to well-correlate with their fibril-formation efficiencies [15]. We checked theoretical propensity to random-coil structure (Pc) by the same algorithm and found that there were unequivocal differences between the three in the region from 132-136, which has a peak in Pc-graph (Fig. 3B) and a trough in the Pβ-graph (Fig. 3A). Suspecting potential significance of the Pc values for cross-seeding efficiencies, we also reviewed a series of cross-seeding experiments of an amyloidogenic peptide derived from tau protein (R3) and its variants, e.g. C-terminally-truncated variants and substitution variants with the serine at the residue 316 replaced with either proline (R3-S316P) or alanine (R3-S316A) [16]. In the experiments, fibril formation of R3-316P was very slow on its own but could be enhanced by adding fibrils of wild-type R3 or other variants as a seed. However, R3-S316A and C-terminally-truncated variants which lack the serine at 316 did not exert the seeding effects. On Pβ-graphs of the peptides, R3-S316A and R3-S316P had peaks at different positions (Fig. 3C, **blue and red arrows**). On the Pc-graphs, interestingly R3-S316P and the C-terminally-truncated variants which could seed R3-S316P, namely ΔCR3SK and ΔCR3S, showed similar curves whereas another truncated variant ΔCR3 which could not seed showed a pattern closer to wild-type R3 (Fig. 3D). The possible relation between the seeding effects and Pc is discussed in detail below.

**Figure 3.**
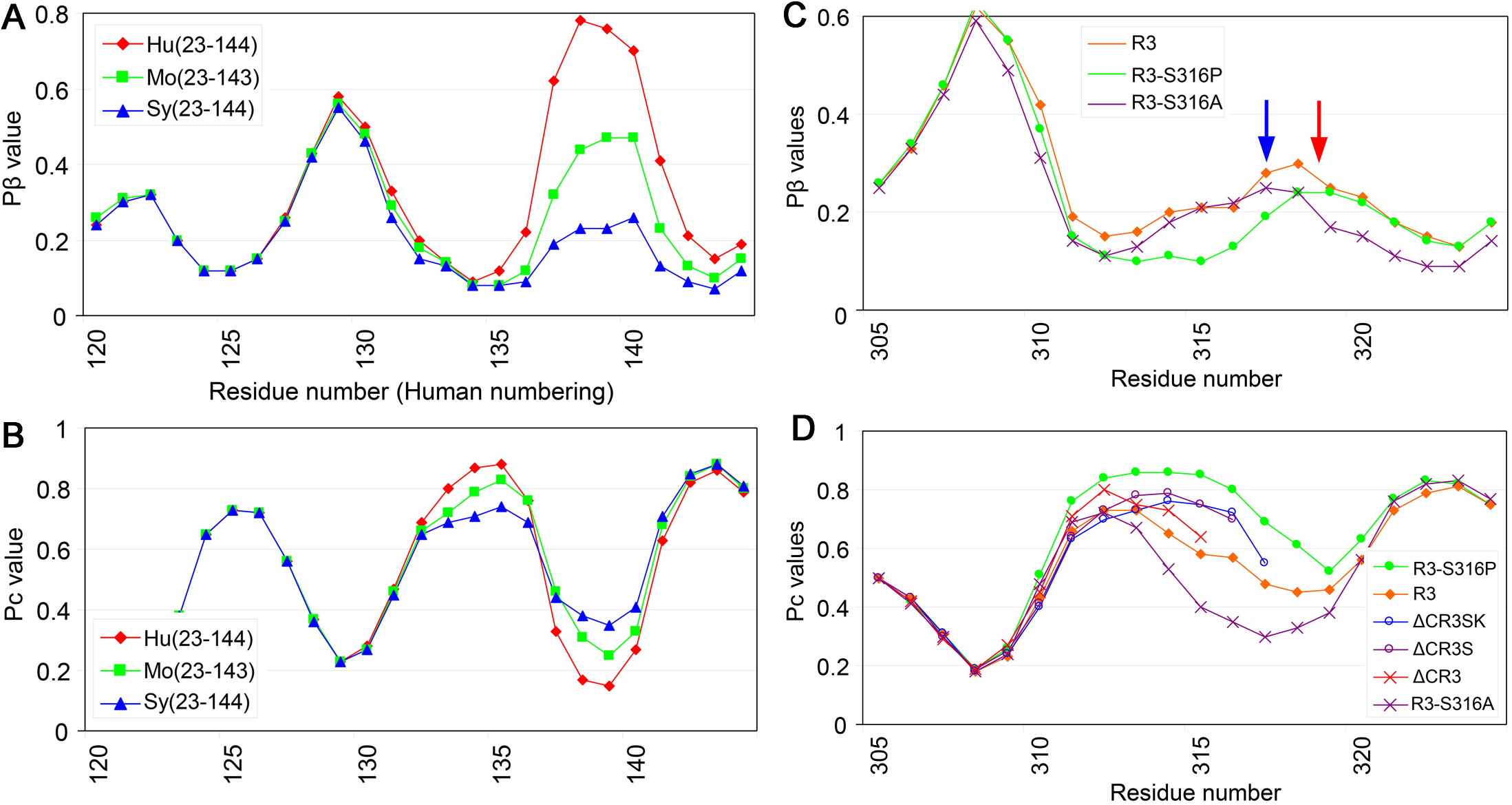
Smaller discrepancies in Pc values in P(3-trough regions are advantageous for cross­ seeding of amyloids. **A.** Pβ-graphs of Hu(23-144) and C-terminally-truncated mouse and Syrian hamster PrP, Mo(23-143) and Sy(23-144), respectively, which are equivalent to Hu(23-144) [15). Note that Hu(23-140) is the highest in the peak at ~140, while Sy(23-144) is the lowest and Mo(23-143) positioned in between. Interestingly, the heights of the peak seem to correlate with their amyloid formation efficiencies: Hu(23-144) most efficient, Sy(23-144) least efficient and Mo(23-143) intermediate. **B.** Pc-graphs of Hu(23-144), Mo(23-143) and Sy(23-144). Note that the Pc-peak at ~135 is located at a position between two P -peaks, P -trough (compare with **Fig. 3A)** and the Pc-peak is highest in Hu(23-144), lowest in Sy(23-144) and intermediate in Mo(23-143). **C.** Pβ-graphs of tau-derived amyloidogenic peptide, R3, and its substitution variants, R3-S316A and R3-S316P, whose serine residue at the codon 316 was replaced with alanine or proline, respectively [16). Note that the two variants have P peaks at different positions (red and blue arrows). The wide P trough of R3-S316P might explain the inefficient fibril formation due to too high freedom of motion in the loop/kink between the two strands. **D.** Pc-graphs of R3, the two substitution variants, and C-terminally-truncated variants, b.CR3SK, b.CR3S and b.CR3. b.CR3S and b.CR3SK have the serine at the codon 316, whereas b.CR3 lacks it. Note that the curves of b.CR3S and b.CR3SK are similar to R3-S316P but the curve of b.CR3 is closer to the wild-type R3. R3-S316A is rather far from R3-S316P.

### Testing biological relevance: Effects of a prion-resistant polymorphism

We analyzed a conversion-resistant polymorphisms of human PrP, valine at the residue 127 (V127) [17], in comparison with another polymorphism, valine at 129 (V129). V127 showed a shift of the Pβ peak at 129 (Fig. 4A, **red arrow**) with elevation of the region 125-128 (**black upward arrows**) and sink of the peak at 122 (**black down-ward arrow**), while V129 simply elevated the peak at 129 despite they are only single-residue apart (Fig. 4A, **blue arrow**). Comparison of Pc-graphs of V129 and V127 (Fig. 4B) revealed that the Pc-peak of V127 in the Pβ-trough region (**gray curve**) were rather lower and narrower than that of the wild-type or V129. Possibly, the remodeling of Pβ and Pc by V127 is responsible for the resistance to the conversion to PrP^Sc^, because a PrP with a shifted β-strand would have difficulty in arrangement with the counterpart of the template PrP^Sc^ side-by-side to form stable parallel-in-register β-sheets. The same holds true to the tau-derived peptides R3-S316A and R3-S316P, which had Pβ peaks at different positions and did not cross-seed (Fig. 3A). Besides, properties of the loop/kink regions connecting the β-strands might affect the parallel in-register β-sheet formation as well (Fig. 4C). Hennetin and colleagues suggested significance of β strand-loop-β strand motif (β-arch), especially of the loop region (β-arc), for parallel in-register β-sheet formation [18]. They created an algorithm “ArchCandy” for prediction of amyloidogenicity based on their hypothesis, which correctly predicted amyloidogenicity of some representative amyloids and even structures of the β-arches [19]. Although ArchCandy failed to explain the results of the aforementioned experiments of PrP23-144, their ideas seem rather reasonable and promising. If the Pc values are another representation of the structures of the β-arcs, the varied cross-seeding efficiencies of PrP23-144 or tau R3-derived peptides are explainable in the light of the β-arc theory, i.e. peptides with similar Pc values in the Pβ-trough can cross-seed because they have similar β-arcs. As another explanation for the varied cross-seeding efficiencies, it is also intuitively conceivable that a flexible loop/kink of the substrate peptide would be more adaptive and facilitate arrangement of the β-strands in appropriate positions and orientations for incorporation into the template, particularly in heterologous reactions (Fig. 4D, **middle left panel**), whereas a rigid loop/kink has difficulty. From this view point, the above cross-seeding efficiencies of PrP(23-144) of human, mouse and Syrian hamster could be explained like this: as the lowest Pc of Sy(23-144) reflects a rigid loop/kink, Sy(23-144) is least adaptive and cannot be cross-seeded, whereas Mo(23-144) or Hu(23-VI. has a more flexible adaptive loop/kink and is prone to cross-seeding in heterologous reactions. On the other hand, a less flexible loop/kink might be advantageous in parallel in-register β-sheet formation in homologous reactions because the restricted freedom of motions would enhance arrangements of β-strands (Fig. 4D, **bottom left panel**). Seeking more quantitative evidence for the biological relevance of Pβ and Pc, we utilized the data of fluorescence intensities of GFP fused to Aβ42 mutants [20] (**Supplementary Results and Discussion** and **Supplementary Fig. S1**). Briefly, the Pβ alone showed a fair correlation with the fluorescence intensity (correlation coefficient -0.768) (**Supplementary Fig. S1B**), an indicator of aggregation tendency, but the ratio of Pβ and Pc exhibited even better correlation (correlation coefficient - 0.833), supporting our view (**Supplementary Fig. S1C**). It should be noted that the view does not necessarily specify the parallel in-register β-sheet model as the correct one, because β-arches are also important structural components of β-solenoids [18]. Although it is much easier to picture the conversion reaction of parallel in-register amyloids, other evidences are required to specify which is correct.

**Figure 4.**
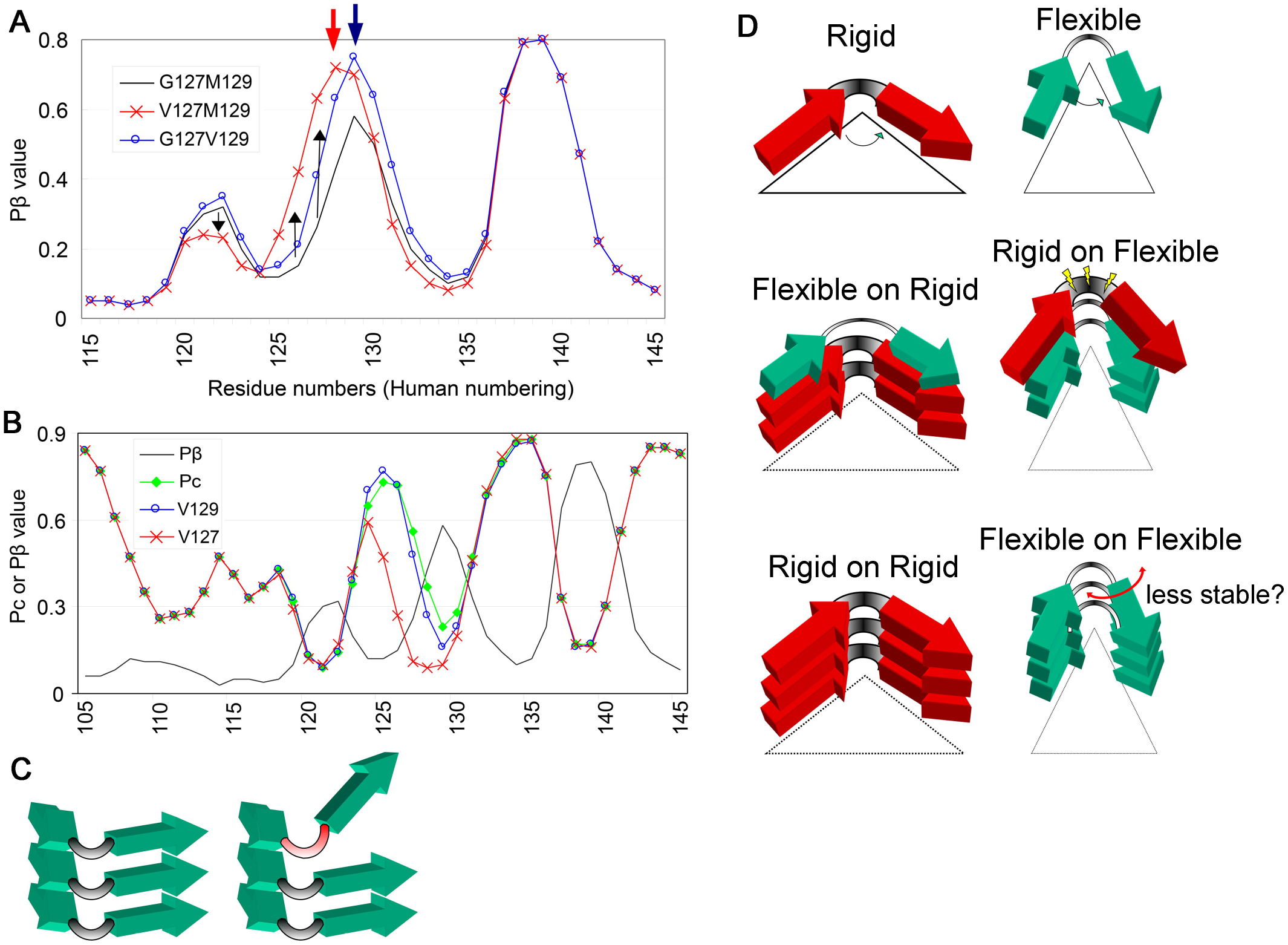
Positions of the β-strands and flexibility of the loop/kink might affect cross-seeding efficiencies. **A.** Anti-prion effects of a polymorphism V127 can be attributable to alterations in P and Pc. Comparison of Pβ-graphs of human PrP with polymorphisms V127 (V127M129), V129 (G127V129) and the wild type PrP (G127M129). The red and blue arrows indicate the crests of peaks of V127M129 and G127V129, respectively. Black arrows indicate changes of Pβ values of V127M129 from those of the wild-type. **B.** Comparison of Pc-graphs of wild-type human PrP and those with the polymorphism V129 or V127. The gray curve represents the Pβ-graph of the wild-type human PrP. **C.** A schematic illustration of a possible influence of the loop/kink region on parallel-in-register -sheet formation. If the properties of loops/kinks, e.g. flexibility or length, are different, the peptides could not form a stable parallel-in-register β-sheet. **D.** Schematic illustrations of two types of β-loop-β motifs (β-arch) with either a rigid (upper panels, red arrows with a thick intervening loop/kink) or a flexible loop (green arrows with a thin intervening loop). The middle panels illustrate heterologous template-substrate conversion reactions: the left panel shows a reaction where the template has the rigid loops/kinks and the substrate has a flexible loop, and the right panel shows the opposite case. The bottom panels illustrate homologous substrate-template conversion reactions.

### Versatility of Pβ in interpretation of transmission results: Introduction of relative ΔPβ

As Pβ seemed to have biological relevance, we assessed versatility of numerical conversion of the primary structure of PrP in analysis of interspecies transmission experiments. However, variation in the heights of the peaks made Pβ-graphs inconvenient for comparison between PrP from different species; for example, it obscures significance of differences in Pβ in regions with relatively-low Pβ values. Considering that PrP^Sc^ already has β-sheets even in such regions irrespective of theoretical Pβ and that refolding of the substrate PrP^C^ is assisted by the β-sheet-rich PrP^Sc^ in template-guided manner, we reasoned that a parameter relative to the absolute Pβ values of the template PrP would more appropriately represent impacts of the differences in the primary structures. Therefore, we adopted a parameter, relative ΔPβ, which represents deviations in Pβ between the substrate PrP^C^ and the template PrP^Sc^, cf. Material and Methods. We applied the new parameter to PrP of three intensively-investigated species, mouse, bank vole and Syrian hamster and found it useful for comparison between different species (Fig. 5A). Note that the mouse and bank vole PrP were especially different in the region 150-175, which comprises the two residues reported to be responsible for the unique transmission properties of bank vole, namely 154 and 169 [21]. As expected from the formula, the ΔPβ-graph patterns were greatly changed depending on the reference species, e.g. human (Fig. 5B) or elk (Fig. 5C).

**Figure 5.**
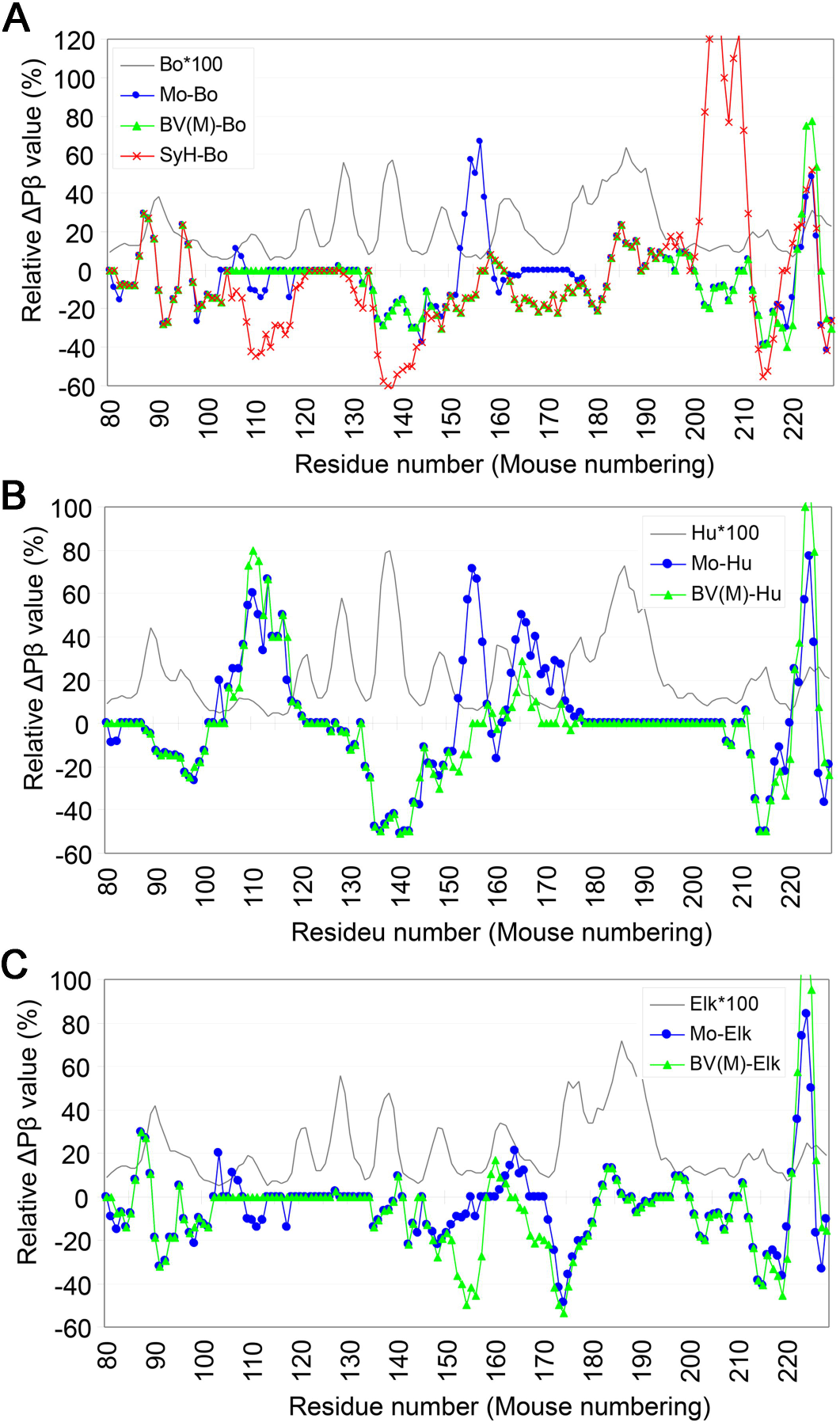
Relative. aPβ-graphs can reveal subtle differences in PJ3 between PrP from different species. **A** Comparison of Pβ-graphs of PrP from mouse (Mo), Syrian hamster (SyH) and bank vole with methionine at the polymorphic codon 109 [BV(M)] relative to bovine PrP (Bo). The gray curve without any marker represents a PJ3-graph of bovine PrP, multiplied by 100 to enable comparison with the Pβ-graphs (Bo*100). **B and C.** Comparisons of Pβ-graphs of mouse and bank vole with M109 relative to human PrP **(B)** and elk PrP **(C).** Note that the relative PJ3-graph patterns of are substantially changed depending on what species is used for the reference.

### Interspecies transmissions between various hamsters

Transmission efficiency of a prion in interspecies transmissions is hypothesized to depend on whether the conformation of the inoculated PrP^Sc^ is included within the repertoire of conformations that PrP of the recipient species can potentially adopt [22]. As an indicator of deviations in Pβ between two PrP, we thought relative ΔPβ-graph could reflect differences in the repertoire of conformations between substrate and template PrP. We analyzed experimental transmissions of Syrian hamster prion Sc237 to Chinese hamster and Armenian hamster [23] and another series of transmission of Syrian hamster prion 263K, which is purported to be the same as Sc237, to a similar set of hamster species [24]. Inoculations of Sc237 to Armenian hamster or Chinese hamster developed diseases after incubation periods of ~174 days or ~344 days, respectively [23]. As the primary structures of PrP of the two hamster species are distinct at three residues, specifically 102, 107 and 111, the difference in the incubation periods seemed to be due to the residues. On ΔPβ-graphs relative to Syrian hamster (Fig. 6A), the differences in the primary structures were recognized as distinct sizes of positive peaks in the region 95-120 and the sizes seemed to correlate with their incubation lengths: the higher and wider the positive ΔPβ peak centered at 110, the longer the incubation. This implied that Sc237 disfavors high Pβ values in the region, exemplifying that high Pβ is not necessarily advantageous for efficient transmission. The long incubations of ~314 days in the transmission of Armenian hamster-passaged Sc237 to Chinese hamster, whose relative ΔPβ showed a large positive peak in the region 110-120 (Fig. 6B, **red curve**), also supported the view. In the back-transmissions of Chinese hamster-or Armenian hamster-passaged Sc237 to Syrian hamster, interestingly, incubation periods were relatively short and not very different, ~121 days or ~113 days, respectively. Although these phenomena were regarded as the evidence for host factors determining the incubation length [23,24], these seemed explainable also by the influences of properties of loops/kinks on PrP^C^-PrP^Sc^ conversion.

**Figure 6.**
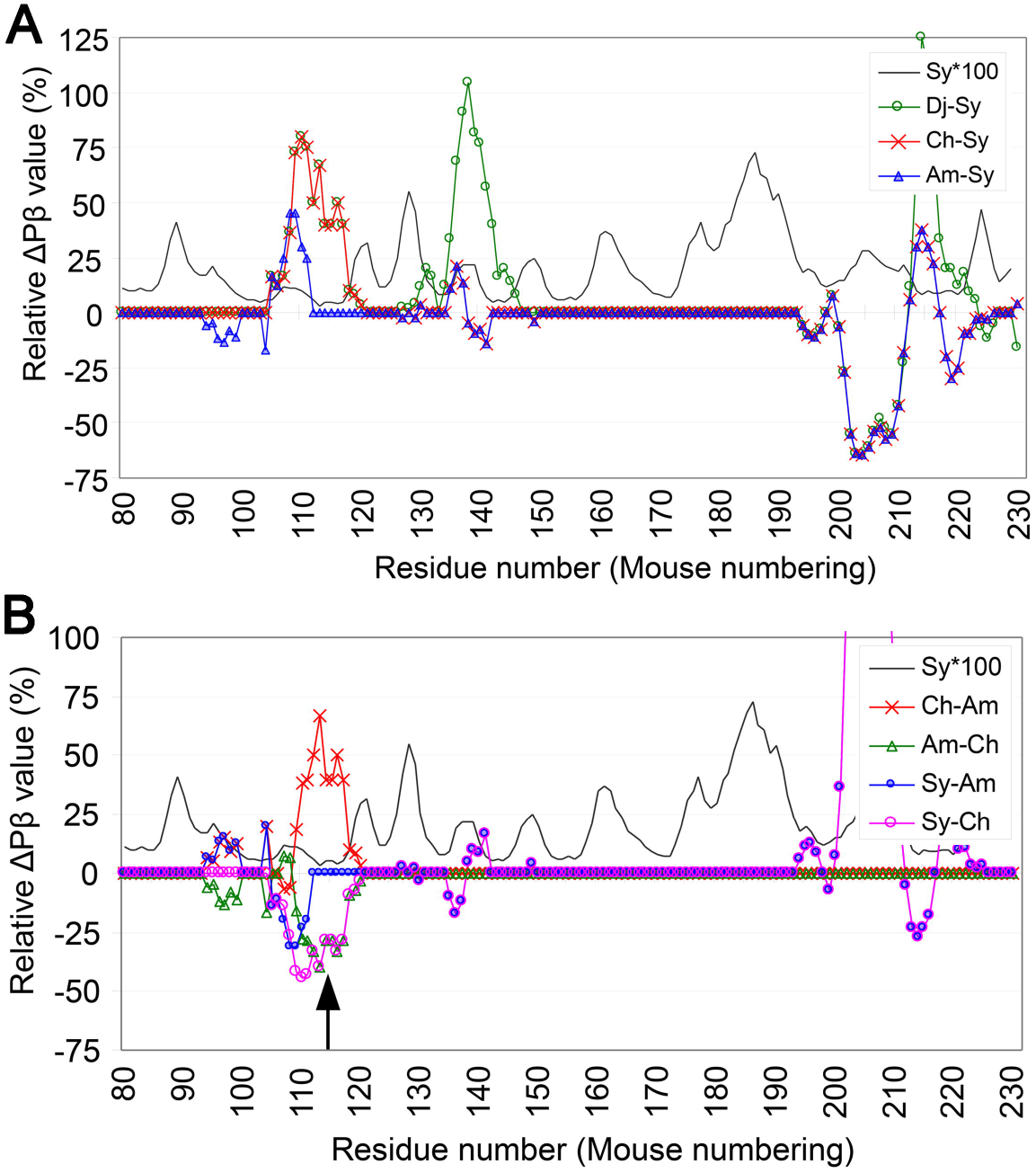
Relative ΔΡβ-graphs of transmissions of S0237 among various hamster species visually aid interpretation of the results. **A.** ΔΡβ-graphs of various hamsters relative to Syrian hamster PrP, for transmissions of S0237 from Syrian hamster to the other hamster species. The reference species are the donors of the transmissions and the former species are the recipients. Sy, Syrian hamster. Ch, Chinese hamster. Am, Armenian hamster. Dj, Djungarian hamster. **B.** ΔΡβ-graphs for back-transmissions of Sc237 from the different hamster species to Syrian hamster, and for the transmission between Armenian and Chinese hamsters. The upward arrow indicates the position of the Pß-trough at the residue ~115.

The relative ΔPβ-graphs of the back transmissions were almost mirror images of the ΔPβ-graphs of the forward transmissions, with the positive peaks in the latter inverted to negative peaks in the former and *vice versa* (Fig. 6B). In the transmission from Chinese hamster to Syrian hamster, a substantial part of the negative peak at the region 105-120 matches the Pβ-trough (Fig. 6B, **arrow**). If a Pβ-trough represents a loop/kink as discussed above, lower Pβ in the trough could represent a more flexible loop/kink and more adaptive to a heterologous template (Fig. 4D); that might be the reason for the relatively small difference in incubation periods. Notably, Djungarian hamsters showed relatively short incubation periods for Syrian hamster 263K, despite the same ΔPβ-graph pattern as Chinese hamster in the region 95-120 (Fig. 6A, **green curve**), as if the negative effects of the positive peak at 110 was neutralized by another positive peak at ~140. As one possibility, this is attributable to stronger interactions of Djungarian-hamster PrP with the template PrP^Sc^ through the interface at ~140 because of the higher Pβ. The strong interactions could keep substrate and template PrP molecules close long enough for the other interface region at ~110 to complete structural changes. Another possibility is discussed below.

### Change of the host range of C-BSE through interspecies transmission

TSEs often drastically change transmission properties after interspecies transmissions. For example, sheep-passaged C-BSE is more efficiently transmissible to transgenic (Tg) mice expressing human PrP [25], elk PrP [26] or porcine PrP [27]. Mouse-passaged C-BSE becomes transmissible to Syrian hamsters [28]. Ferret-adapted CWD more efficiently infect Syrian hamsters [29]. Not only enhancement of virulence, loss of transmissibility to the original species can also occur, as seen in the poor transmission of the CWD passaged in bank vole-PrP-expressing Tg mice to elk PrP-expressing Tg mice [30]. We expected that relative ΔPβ analysis could visually aid interpretation of the results of those interspecies transmissions and provide a clue to the underlying mechanisms. The relative ΔPβ-graph of the transmission of sheep-passaged C-BSE to elk-PrP-expressing Tg mice (Fig. 7A, **green curve**) apparently showed less deviation from the base line than the ΔPβ-graph of the transmission of cattle C-BSE to elk (Fig. 7A, **red curve**), suggesting a possibility that the enhanced transmissibility might be due to improvement of the deviations in Pβ between the template PrP^Sc^ and the substrate PrP^C^. Relative ΔPβ-graphs of cattle-ovine-porcine serial transmission of C-BSE also showed the improvement of deviations in the regions 150-165 and 180-195 (Fig. 7B, **bottom brackets**). Possibly, the improvement of deviations between the template PrP^Sc^ and the substrate PrP^C^ in those regions might compensate for the remaining deviation in the intervening region.

**Figure 7.**
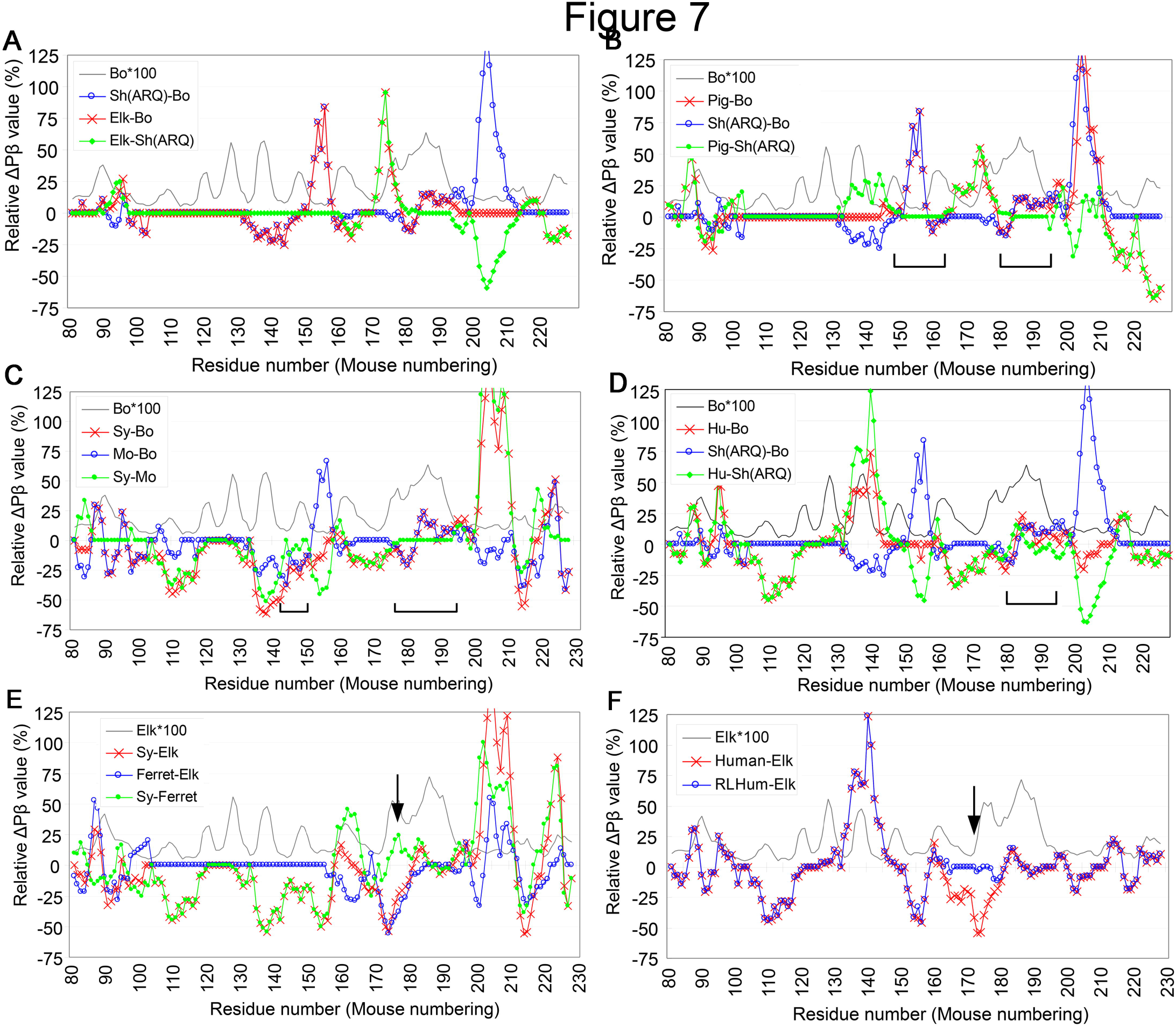
Relative ΔΡβ-graphs of interspecies transmissions visually aid interpretation and are suggestive of how changes of host ranges could occur. **A.** Relative ΔΡβ-graphs of transmissions of C-BSE from cattle to sheep with ARQ polymorphism [Sh(ARQ)-B0], from cattle to Tg mice expressing elk PrP (Elk-Bo) and transmission of ovine PrP-adapted C-BSE to Tg mice expressing elk PrP [Elk-Sh(ARQ)][26], The latter species are donors and the former species are recipient of the transmissions. The blue, red and green curves represent the efficient, the inefficient and the improved transmissions, respectively. **B.** Relative ΔΡβ-graphs of transmissions of C-BSE from cattle to Tg mice expressing porcine PrP (Pig-Bo) and of ovine PrP-adapted C-BSE to Tg mice expressing porcine PrP [Pig-Sh(ARQ)]. The bottom brackets indicate regions which were flattened by the passage through the sheep [27], **C.** Relative ΔΡβ-graphs of transmissions of C-BSE from cattle to mouse (Mo-Bo), from cattle to Syrian hamster (Sy-Bo) and mouse-adapted C-BSE to Syrian hamster (Sy-Mo) [28], **D.** Relative ΔΡβ-graphs of transmissions of C-BSE from cattle to Tg mice expressing human PrP (Hu-Βο) and of ovine PrP-adapted C-BSE to human [Hu-Sh(ARQ)] [25], **E.** Relative ΔΡβ-graphs of transmissions of CWD from elk to ferret (Ferret-Elk), from elk to Syrian hamster (Sy-Elk) and ferret-adapted CWD to Syrian hamster (Sy-Ferret). Note that the large negative peak at around the residue 175 disappeared after passage through ferret (arrow) [29], Relative ΔΡβ-graphs of transmissions of CWD from elk to Tg mice expressing human PrP (Human-Elk) and from elk to Tg mice expressing human PrP with elk residues in the region from the residue 166 to 174 (RLHuman-Elk) [31], The arrow indicates the region where the graph pattern was changed by the substitutions.

Although less apparent than those two cases, cattle-mouse-Syrian hamster serial transmission of C-BSE would be also explained in the same manner: The ΔPβ-graph is flattened in the region 175-195 and the deviation in the region 145-150 is also improved approaching to the baseline (Fig. 7C, **bottom brackets**). Unlike the foregoing cases, in the transmission of sheep-adapted BSE to Tg mice expressing human PrP, the relative ΔPβ-graph did not become flat in the region 180-195 but the deviation in the sheep-to-human transmission seemed improved compared with that of the cattle-to-human transmission (Fig. 7D).

Collectively, those results imply that the improvements in the deviations in the regions 150-165 and 180-195 might facilitate transmissions of C-BSE to originally less-susceptible species. This implication requires validation by other types of experiments. If proven true, it strongly supports the legitimacy of our approach.

### Change of the host range of CWD through interspecies transmission

In the relative ΔPβ-graph of transmission of ferret-passaged CWD to Syrian hamster (Fig. 7E, **green curve**), one positive peak newly appeared in the region 170-180, which has the crest at the similar position as the Pβ peak at ~175 of elk PrP (Fig.7E, **arrow**), compared with transmission of elk CWD to Syrian hamster (**red curve**). This seemed reminiscent of a ΔPβ-graph of transmission of CWD to Tg mice expressing chimeric human PrP with elk residues in the region 166-174, which greatly facilitated transmission of CWD, compared with Tg mice expressing pure human PrP [31]. On relative ΔPβ-graph, the negative peak in the region 160-180 seen in the ΔPβ-graph of elk-to-human transmission was completely buried in the chimeric PrP (Fig. 7F, **arrow**). Those findings suggest that smaller discrepancies in Pβ in the region between substrate and template are advantageous for propagation of CWD prion.

### Mechanism of strain diversity from our view point

Relative ΔPβ is unambiguously defined by the primary structures of PrP of donor and recipient species. On the other hand, even between the same pair of species, some prion strains are transmissible, whereas others are not. For example, cattle C-BSE is transmissible to mouse but L-BSE is not. This could be explained by strain-specific patterns of usages and/or predominance of certain interfaces among the multiple interfaces of PrP [12]: some strains favor high Pβ values in a certain interface region, while other strains demand low Pβ in the same region. Such variations could presumably stem from either stochastic events, presence of cofactors and/or environments during the initial nucleation of PrP^Sc^, and be inherited to the offsprings.

### Problems to be addressed in the future: intrinsic properties

Beside Pβ values, other attributes of β-strands of the interface regions also theoretically affect interaction efficiencies between the substrate PrP^C^ and template PrP^Sc^, including interactions through side chains (e.g. steric zipper or electrostatic interactions), twist/tortion of β-strands, and positioning and orientations relative to the other strands [32]. To address those factors, molecular dynamic simulation of parallel-in-register amyloids would be necessary for future studies. Apart from that, an algorithm which combines the characteristics of ArchCandy and the secondary-structure prediction seems feasible because both of them depend solely on the primary structure for calculation. Such an algorithm might further improve the prediction of regional structures of PrP^Sc^, while providing an insight into another aspect of intrinsic propensities of PrP.

### Problems to be addressed in the future: extrinsic factors

We demonstrated that Pβ and Pc carry a substantial information about regional structures of amyloids. On the other hand, we are aware that there are many other factors which definitely affect conversion efficiency of the substrate PrP^C^ but are incalculable by the algorithm, e.g. post-translational modifications like GPI anchor, N-linked glycans and the disulfide bond between the second and the third helices. Particularly, the disulfide bond can cause unpredictable results by keeping high-Pβ regions in vicinity and precipitating their interactions. Even without a disulfide bond, if PrP^Sc^ is compactly folded as the parallel in-register β-sheet model illustrates [8], interactions between high-Pβ regions can occur in theory: Possibly the aforementioned relatively-short incubations in transmissions of 263K to Djungarian hamster could be a manifestation of such effects. Certain types of phospholipids or nucleotides which are postulated to be “cofactors” for PrP^C^-PrP^Sc^ conversion might also affect actual β-sheet propensities [33–35]. Those extrinsic factors should greatly contribute to the behaviors and the strain diversity of prions and always have to be considered in investigations with Pβ and Pc.

By addressing those problems, our approach can lead to development of new investigation tools which are applicable to not only prion but also other amyloids, e.g. Aβ or α-synuclein. Advances of understanding of proptein-protein interactions underlying amyloid propagation and cross-seeding can consequently benefit even protein science/engineering as well as investigation of neurodegenerative diseases.

## Materials & Methods

### Primary structures of PrP from various species

Amino acid sequences of PrP of various species were obtained from the website of UniProt (http://www.uniprot.org/). The species studied here were mouse (*Mus musculus*), Syrian hamster (*Mesocricetus auratus*), bank vole (*Myodes glareolus*), human (*Homo sapiens*), elk (*Cervus elaphus nelsoni*), cattle (*Bos taurus*), sheep (*Ovis aries*), pig (*Sus scrofa*), Chinese hamster (*Cricetulus griseus*), Armenian hamster (*Cricetulus migratorius*), Djungarian hamster (*Phodopus sungorus*) and ferret (*Mustela putorius furo*). A region from the tryptophan of the second-last repeat of the octapeptide repeat region (OPR), e.g. the 80th residue of mouse PrP, to the putative GPI-attachment site (e.g. the 231st residue of mouse PrP) was excised and rendered to the secondary structure prediction by neural network. All the residue numbers are in mouse numbering unless otherwise noted.

### Secondary structure prediction

Neural network prediction method [11] available at the website of the Center for Information Biology, Ochanomizu University (http://cib.cf.ocha.ac.jp/bitool/MIX/) was used for secondary-structure prediction of the selected region of PrP described above. Then, values of the “Neural Network Prediction 2” of the prediction results were adopted for Pβ values of PrP and processed for creation of Pβ-graph and calculations of relative ΔPβ values.

### Definition of Relative Δ**P**β

For acquisition of relative ΔPβ, the selected regions of PrP from different species were arranged so that the proline at the residue 101 of mouse PrP and the corresponding proline residues of the PrP from other species are in the same row of the work sheet, as presented in **Supplementary table 1**. By doing this, most of the corresponging residues of PrP from different species are arranged in the same row.

Relative ΔPβ between PrPs from species A and species B at a given residue is defined by the flowing formula:

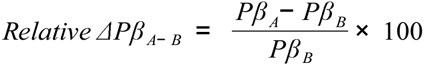

where Pβ_A_ and Pβ_B_ are Pβ value at the corresponding residues of PrP of the species A and B, respectively, and in this case species B is referred to as the “reference species”.

### Correlation analysis of Pβ^max^ or Pβ^max^/Pc^max^ with fluorescence intensities of mutant Aβ42

Pβ^max^ is the maximum value of Pβ of a given Aβ42 mutant in the interval form the residue 16 to 23 (**Supplementary Fig. S1A**). Likewise, Pc^max^ is the maximum value of Pc in the interval from the residue 13 to 19. For the fluorescence intensities, we measured the lengths of the bars of the graph of the relative fluorescence intensities (presumably representing the mean values) in the paper by De Groot et al. [20] and compared with the Pβ^max^ or the ratio between Pβ^max^ and Pc^max^, Pβ^max^/Pc^max^. The scatter plots and the correlation coefficients were generated using the functions in the Excel2013.

## Abbreviations

Aβ42: β-amyloid of 42 residues
BSE: bovine spongiform encephalopathy
CJD: Creutzfeldt-Jakob disease
CWD: chronic wasting disease
DNI: dominant-negative inhibition
GFP: green fluorescence protein
GPI: glycosylphosphatidylinositol
OPR: octapeptide-repeat region of prion protein
Pα: theoretical α-helix propensity
Pβ: theoretical β-sheet propensity
Pc: theoretical random-coil propensity
PrP: prion protein
PrP^C^: normal isoform of PrP
PrP^Sc^: disease-associated conformer of PrP
Tg mouse: transgenic mouse
TSE: transmissible spongiform encephalopathy

## Acknowledgment

We especially appreciate Professor Kei Yura, Ocha-no Mizu University, generously allowing us to use the secondary structure prediction software on his website along with very helpful advices. We thank Professor Kajava for kindly providing us with ArchCandy. We also thank Professor Kuwata and Professor Kitao for critical reading of the manuscript. This study was supported by Takeda Science Foundation.

## Supplementary Results and Discussion

Relations between Pβ and Pc values and aggregation propensity using the data from Aβ42-GFP fusion protein experiments.

To further test the relation between the Pβ values calculated by the secondary structure prediction algorithm and actual aggregation efficiencies of amyloidogenic proteins, we utilized the data of fluorescence intensities of mutant Aβ42-GFP reported by De Groot et al. [20]. In the experiments, they replaced the phenylalanine at the residue 19 of Aβ42 moiety of the fusion protein with nineteen kinds of amino acids and monitored the mutant's aggregation tendencies by the fluorescence from the GFP moiety; since the alterations in Pβ or Pc mostly occur in a single Pβ-peak or a Pβ-trough, this data set was easy to apply to our system. Comparison between the maximum Pβ values of the Pβ peak (**Fig. S1A**) and the fluorescence intensities revealed a fair degree of correlation with a correlation coefficient -0.768 (**Fig. S1B**). On the other hand, we noticed that Pc in the adjacent Pβ-trough region seemingly changed in association with the fluorescence intensity, although Pc itself did not show substantial correlation with the fluorescence intensities. Since we had suspected a role of Pc in amyloid formation, we compared the fluorescence intensities with Pβ^max^/Pc^max^ ratios. As a result, the correlation coefficient got even better, -0.833, than that of Pβ alone, whereas highly-charged residues and polarized residues with relatively large side chains, e.g. glutamine and threonine, still remained as outliers (**Fig. S1C**). This improvement imply a certain level of influence of Pc of the Pβ-trough region on the amyloid formation. Although direct comparison is difficult, the correlation coefficient on the data set seems comparable with the other amyloid-prediction algorithms [36], suggesting that Pβ and Pc carry substantial structural information of amyloids.

**Supplementary Figure S1.**
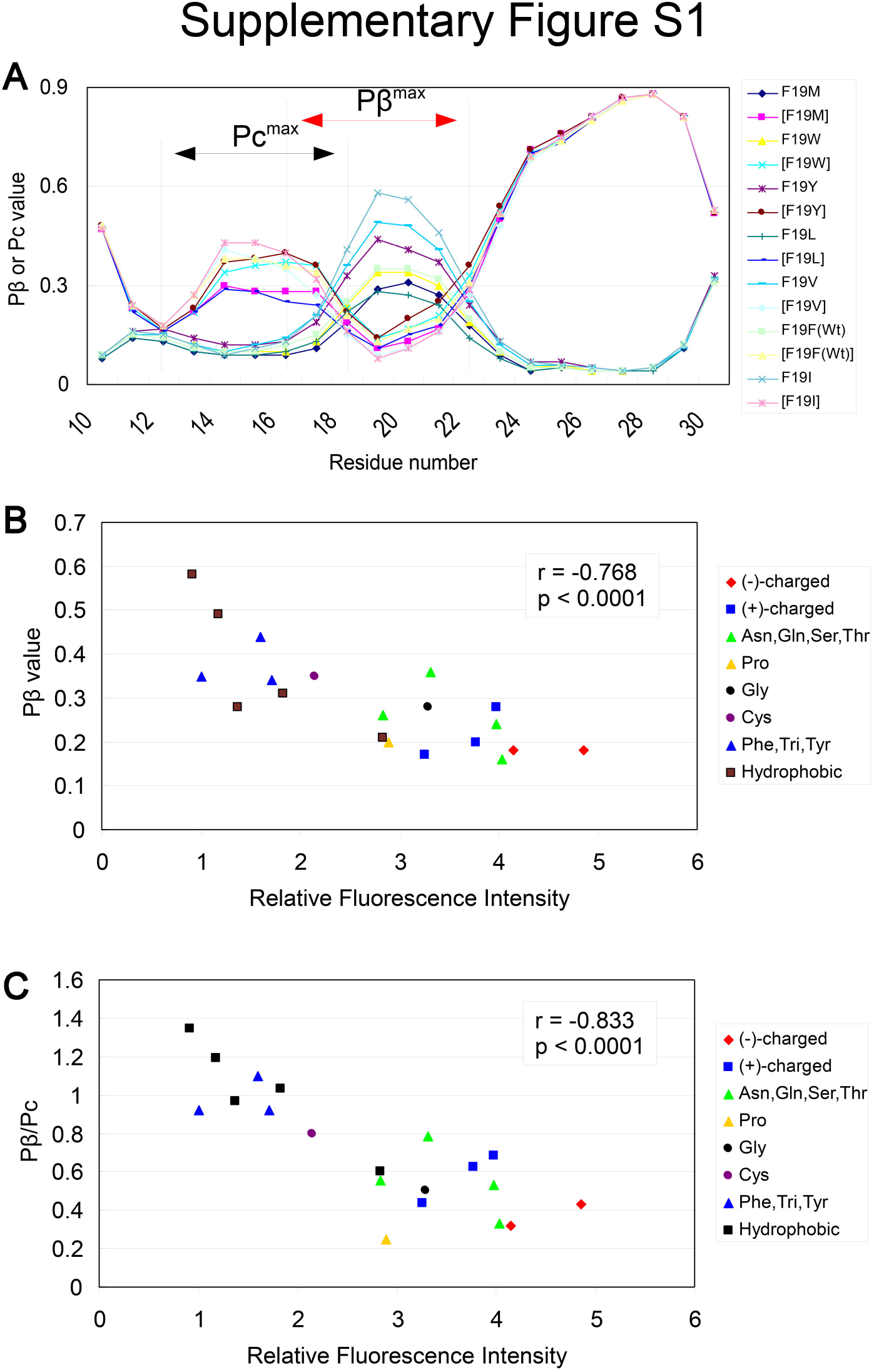
Pß values have a fair correlation with aggregation efficiencies of substitution-mutant Aß42 and pß^max^/p_c_^max^ show even better correlation. **A.** A Examples of Pß- and Pc-graphs of some of the mutant Aß42. Pß^max^, the highest Pß value in the interval from 17 to 23. Pc^max^, the highest Pc value in the interval from 13 to 19. The curves labeled with square brackets, e.g. [F19Y], represent the Pc -graph of the mutant, while those without the square brackets represent Pß-graphs. **B.** A scatter plot illustrating the correlation between pß^max^ values of the mutant Aß42 and relative fluorescence intensities of the Aß42-GFP fusion proteins. **C.** A scatter plot illustrating the correlation between the ratio of Pß^max^ and Pc^max^, pß^max^/p_c_^max^, _0_f the mutant Aß42 and relative fluorescence intensities of the Aß42-GFP fusion proteins. Note that the correlation coefficient was improved compared with the scatter plot in **Fig. S1B.**

The finding that many of the outliers are charged residues would mean that they destabilized the amyloid formation of Aβ42 more than they were supposed to by the secondary structure prediction algorithm. This is quite understandable because repulsion between the same electric charges in short distances occur in parallel in-register β-sheet amyloids like Aβ42, where the same amino acids are positioned side-by-side.

